# Mechanics defines the spatial pattern of compensatory proliferation

**DOI:** 10.1101/2021.07.04.451019

**Authors:** Takumi Kawaue, Ivan Yow, Anh Phuong Le, Yuting Lou, Mavis Loberas, Murat Shagirov, Jacques Prost, Tetsuya Hiraiwa, Benoit Ladoux, Yusuke Toyama

## Abstract

The number of cells in tissues is tightly controlled by cell division and cell death, and misregulation of cell numbers could lead to pathological conditions such as cancer. To maintain cell numbers in a tissue, a cell elimination process named programmed cell death or apoptosis, stimulates the proliferation of neighboring cells. This mechanism is called apoptosis-induced compensatory proliferation, which was originally reported more than 40 years ago. While only a limited number of the neigboring cells need to divide to compensate for apoptotic cell loss, the mechanisms that select cells for undergoing division remain an open question. Here we found that the spatial inhomogeneity in mechanotransduction through a growth-promoting transcription co-activator Yes-associated protein (YAP) in the neighboring tissue, accounts for the inhomogeneity of compensatory proliferation. Such inhomogeneous mechanotransduction arises from the combination of the non-uniform distribution of nuclear size, which is inherent in tissues, and the non-uniform pattern of mechanical force applied to the neighboring cells upon apoptosis. Our findings from a mechanical perspective complement the current biochemical understanding of compensatory growth and provide additional insights into cellular functions of how tissue precisely maintains its homeostasis.

## Introduction

Apoptosis, or programmed cell death, is a mechanism by which unnecessary, aged, or damaged cells are eliminated [1]. To maintain the homeostatic cell number in epithelium and organs, a mechanism by which an apoptotic event induces the proliferation of neighboring cells, called apoptosis-induced compensatory proliferation, takes place. This phenomenon of compensatory proliferation was originally described more than 40 years ago [2]. Haynie and Bryant investigated how *Drosophila* imaginal wing disc responds to damage induced by x-ray irradiation. Although irradiation of *Drosophila* larvae killed ∼60% of the cells in the wing disc, the remaining cells were able to recover and develop as adult wings with normal structure and size. This suggested that there are cellular mechanisms that promote cell division upon tissue damage to compensate the cell loss. Since then, the mechanisms of compensatory proliferation, especially the role of mitogenic signals secreted by the apoptotic cell, have been investigated [3,4]. For instance, in *Drosophila*, it is well characterized that the secretion of Decapentaplegic (Dpp) and Wnt/Wingless (Wg) mitogens from the apoptotic cell results in cell proliferation in neighboring cells [5–9]. In mice [10] and a mammalian cell line [11], the pro-inflammatory metabolite, Prostaglandin E2 (PGE2), is produced and secreted by the apoptotic cell and stimulates cell growth. Notably, it is sufficient for a limited number of the neighboring cells to divide to compensate for the cell loss due to apoptosis. This raises a possibility that biochemical signals are crucial, but may not be solely responsible, to explain this spatial inhomogeneity of compensatory proliferation. More recently, the apoptotic process, especially in the context of cell extrusion, has been associated with mechanical force. An apoptotic cell is expelled from a tissue through the contraction of an actomyosin cable formed in the dying cell as well as in the neighboring cells, or via lamellipodium crawling by adjacent cells [12–16]. This cell extrusion process further alters the surrounding tissue tension and morphogenesis [17–20]. Here we considered the mechanical force associated with an apoptotic process and elucidate how only a small number of cells around an apoptotic cell undergo cell division.

## Results

### Mechanical force propagation and stretching in the neighboring tissue upon apoptosis

To understand how an apoptotic event mechanically influences neighboring cells, we established an *in vitro* platform to quantitatively measure the changes in mechanical force among neighboring tissue upon cell death in Madin-Darby Canine Kidney (MDCK) epithelial cells (Fig. 1A). DNA damage and subsequent apoptotic cell extrusion (Fig. 1B) in the desired cell were induced by using UV laser [14,21] among a tissue that was cultured on deformable substrates made from elastic polydimethylsiloxane (PDMS) gel with fluorescent beads [22]. Upon the induction of apoptosis, a wave-like propagation of bead displacement, which is the proxy for substrate deformation, was observed (Movie 1). Particle image velocimetry (PIV) analysis of the movement of the fluorescent beads (Fig. 1C) showed that initially, the beads moved away from the apoptotic cell (red, t=5min in Fig. 1C) and this displacement was largest at the nearest neighboring cells (Fig. 1H, Methods). In addition to the outward movement, the inward movement of the substrate emerged at the region close to the extrusion site (blue, Fig. 1C). To quantitatively understand the substrate deformation, we plotted the kymograph of the average radial velocity of the beads as a function of distance from the apoptotic cell (Fig. 1D, Methods). In-between the outwardly and inwardly moving substrate, there was a small region of substrate that showed transiently static behavior (i.e. no or balanced inward and outward movement), and this transiently static region shifted away from the extrusion site over time (white, Fig. 1D). We adopted this static region as one of the hallmarks of substrate deformation and found that the wave propagated a distance of 21.3±2.3μm (mean±s.e.m.) away from the dead cells in an hour (Fig. 1G). To understand the cause of the substrate deformation, we treated the tissue with the Rac1 inhibitor NSC23766 to inhibit lamellipodial cell crawling. Lamellipodia crawling is known to occur in neighboring cells and contributes to apoptotic cell extrusion [14,16] and to deform the substrate to the direction opposite to the cell migration. Although we predicted that NSC23755 treatment would diminish substrate deformation upon apoptosis, we found that the wave propagation increased to a distance of 42.8±2.4μm (mean±s.e.m.) in an hour (Fig. 1E-G, Movie 2). We then tracked the tissue dynamics by differential interference contrast (DIC) imaging and found that the cells moved away from the dead cell in both control and NSC23766-treated tissue (Fig. 1I-J, Movie 3). We reasoned that the release of epithelial tissue pre-tension, which is an inherent tissue characteristic generated by actomyosin-mediated contractility, leads to the relaxation and the outward movement of the tissue around the apoptotic cell. Indeed, we previously reported a reduction of the adherens junction molecule E-cadherin between apoptotic and neighboring cells after caspase-3 activation in the apoptotic cell in *Drosophila* epithelia [13] and MDCK cells [23]. This reduction of E-cadherin leads to disengagement of the cell-cell junction between apoptotic and neighboring cells, and a release of the tissue pre-tension [13]. To further validate our reasoning, we compared the tissue pre-tension between control and NSC23766-treated monolayer by laser ablation at the cell-cell junction [24] without induction of apoptosis. The junctional tension in the NSC23766-treated tissue is higher than that of the control tissue (Fig. S1A-C), which is consistent with the larger wave propagation in NSC23766-treated tissue. We further found that the relative position between the focal adhesions of the neighboring cells and the beads embedded in the substrate did not change drastically during wave propagation, indicating there is minimal sliding between tissue and the substrate during wave propagation (Fig. S1D). Together, our data reveals that the tissue moves away from the dead cell due to the relaxation of the tissue pre-tension upon apoptosis. The wave-like dynamics of the substrate gel caused by the release of the pre-tension of tissue upon apoptosis can be explained by a physical model based on a linear elasticity theory [25].

**Figure 1.**
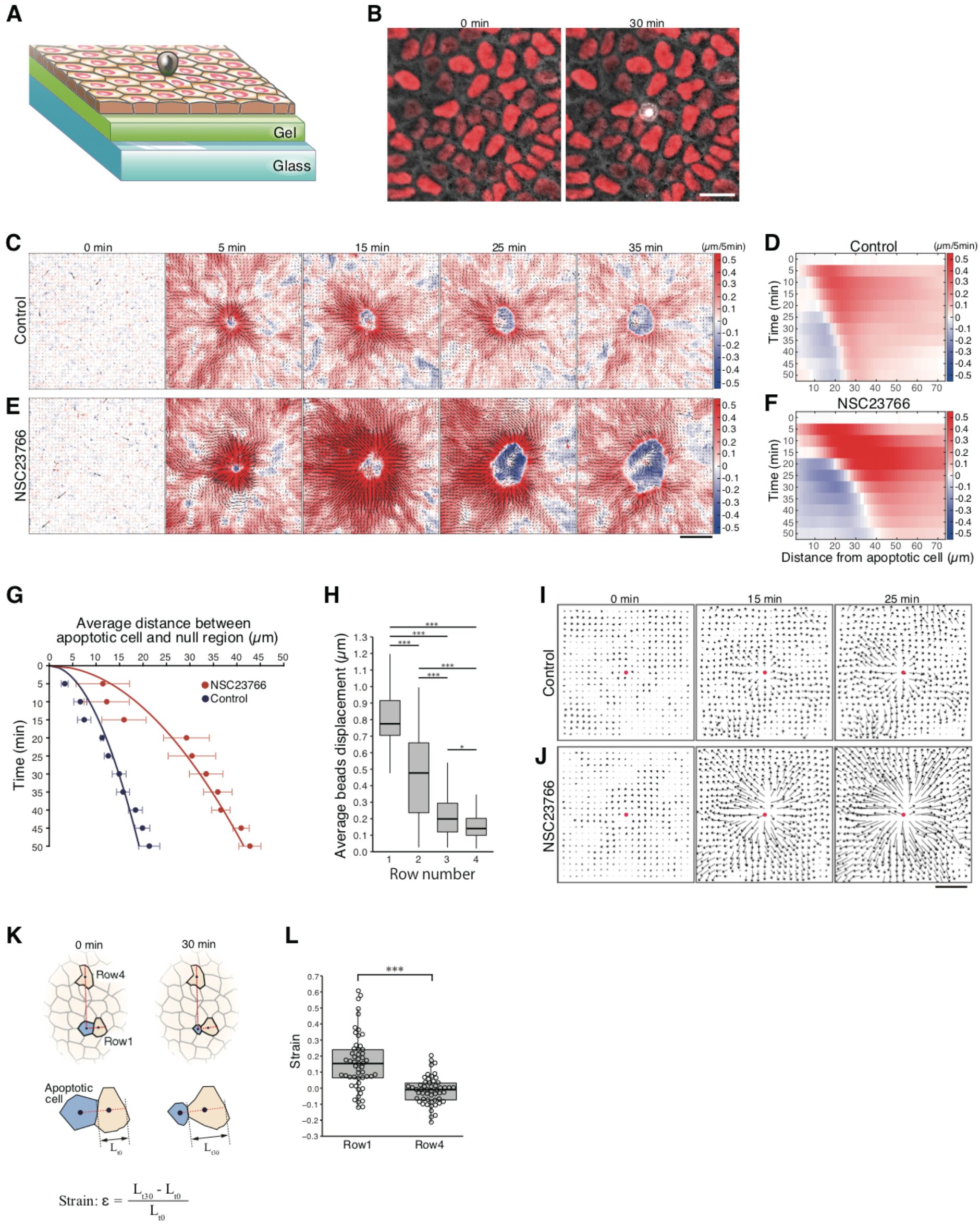
**(A)** Diagram of a confluent MDCK monolayer tissue on the gel-coated glass dish. See Methods. Grey, apoptotic extruding cell; green, silicone gel coated with fibronectin, which overlays on the glass bottom dish. **(B)** Representative snapshots of apoptotic cell extrusion within an MDCK monolayer. (**C** and **E**) Representative time series showing heat map with vectors of radial beads displacement after laser induction of apoptosis for control and NSC23766 treated condition. The color bar indicates the magnitude of the radial outward (red) and inward (blue) displacements in µm. (**D** and **F**) Kymograph of average radial beads displacement in **C** and **E**. **(G)** Average distance from the center of apoptotic cell to the static region where is the boundary between outward and inward vectors. Time 0 min represents the time at laser induction of apoptosis. n = 5 ROIs (control), n = 5 ROIs (NSC23766). **(H)** Average beads displacement 1 min after laser induction for each row from the apoptotic cell. Data are mean ± s.d.; *P < 0.05 and ***P < 0.001, one-way ANOVA and Levene test followed by Tukey–Kramer test. n = 7 ROIs in 3 independent experiments. (**I** and **J**) Representative velocity field from DIC images of monolayers after laser induction in control and NSC23766 treated condition. Red dot, laser irradiation point; length of vectors, proportional to their magnitude. **(K)** Diagram illustrating measurement of cellular strain (ε) for the cells near and far (row1 and row4) from the apoptotic cell (blue). L_t0_ and L_t30_ represent the length of cells 0 and 30 min after apoptosis. See methods. **(L)** Strain of cells in row1 and row4. Data are mean ± s.d.; ***P < 0.001, Mann–Whitney *U*-test. n = 53 cells (row1) and n = 53 cells (row4) from 7 ROIs in 3 independent experiments. Scale bars, 20 µm in **B**; 50 µm in **C, E, I**, and **J**.

A kymograph of the radial average velocity of the beads showed that there was inward movement of the substrate toward the apoptotic cell (blue in Fig. 1C-F). We speculate that this is, in part, due to the formation and the contraction of the actomyosin cable in the neighboring cells [12,16,23]. The appearance of the inward movement emerged around 5∼10min after the initial outward movement, consistent with the timing of formation and initiation of actomyosin cable contraction. These observations imply that the neighboring cells are stretched by apoptotic cell extrusion. Indeed, measuring the strain of the neighboring cells showed that the nearest neighbor cells underwent 16.2±2.3% (mean±s.e.m.) stretch, which is significantly larger than cells further away from the dead cell (Fig. 1K-L). Taken together, we conclude that the cells in the vicinity of the apoptotic cell experience a tensile stretching, which is due to the combination of tissue relaxation and apoptosis-associated actomyosin cable contraction.

### A small number of neighboring cells show nuclear translocation of YAP and cell cycle progression upon apoptosis

To address whether this stretching of the neighboring tissue would further influence the biochemical signaling within the surrounding cells, we examined the dynamics of Yes-associated protein (YAP), a growth-promoting transcription co-activator and an effector of the Hippo pathway [26]. YAP is also known as a mechanotransducer which transforms the physical stimuli that cells experience through mechanosensing into intracellular biochemical signals [27]. Upon mechanical stimuli, YAP translocates into the nucleus, which further promotes downstream transcriptional programs including cell proliferation [28]. We imaged the localization of YAP in the neighboring cells of an apoptotic cell by using MDCK cells that stably express YAP-GFP (Methods), and found that only a fraction of neighboring cells exhibited nuclear translocation of YAP (Fig. 2A-B, Movie 4). To further understand the consequence of apoptosis-associated YAP nuclear translocation, we imaged the cell cycle progression of the neighboring cells by using MDCK cells with the cell cycle reporter Fluorescent Ubiquitination-based Cell Cycle Indicator, FUCCI [29]. The cells were first serum-starved for 24 hours to synchronize the cell cycle at G1 phase. Prior to imaging, serum-free media was replaced with fresh media containing serum. We found that only a few neighboring cells underwent cell cycle progression from G1 to S phase at around 6-7 hours after apoptosis (Fig. 2C-E, Movie 5). The nearest neighbor cells and next-nearest neighbor cells to the apoptotic cell had a higher probability of mitosis, compared to cells further away from the dying cell (Fig. 2F-G) and underwent mitosis at around 16-18 hours after apoptosis (Fig. 2H). Such characteristics are distinct from the cells irrelevant to and further away from apoptosis (Fig. 2F-H). Our data consistently showed that only a few neighboring cells undergo cell division upon apoptosis.

**Figure 2.**
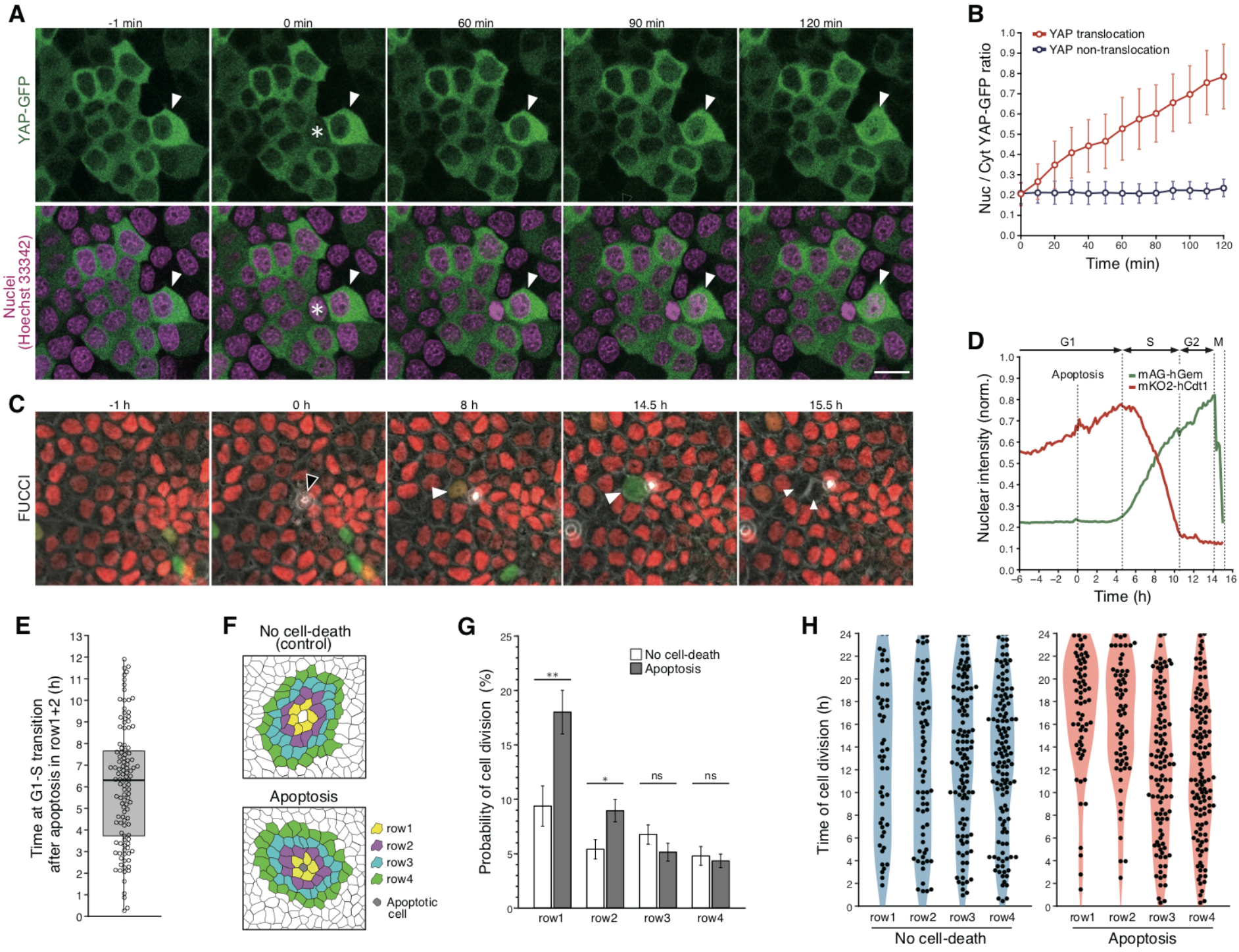
**(A)** Representative time series showing YAP-GFP nuclear translocation (white arrowhead) within an MDCK YAP-GFP monolayer. Time 0 min represents the time at laser induction of apoptosis (asterisk). **(B)** Nuclear/cytosolic (Nuc/Cyt) YAP-GFP ratio in cells surrounding apoptotic cell after laser induction. Cells were classified as YAP translocation (I_nuc/cyt_ ≥ 0.5) or non-translocation (I_nuc/cyt_ < 0.5) 2 h after laser induction (see Methods). n = 42 cells in 10 images from 6 independent experiments. **(C)** Representative time series showing a cell proliferation (white arrowhead) after apoptotic cell extrusion (black arrowhead) within an MDCK FUCCI monolayer. **(D)** Normalized (norm.) fluorescence intensities of the mKO2-Cdt1 (red) and mAG-hGem (green) in a cell as shown in **C** (cyan arrowhead). **(E)** The distribution of time at G1-S phase transition in cells surrounding the apoptotic cell. n = 114 cells from 54 ROIs in 6 independent experiments. **(F)** Illustration depicting segmented cells in ROIs with no dead cells or with an apoptotic cell. Cells in each row (row1-4) surrounding a central living or apoptotic cell are color-coded. **(G)** Probability of cell division for each row within 24 hours after extrusion. Cells in which G1-S phase transition was detected before extrusion were excluded. Data are mean ± s.e.m.; not significant (ns), **P = 0.0013 and *P = 0.0157, Mann–Whitney *U*-test. **(H)** The distribution of cell division time for each row. Time 0 h represents the time at apoptotic cell extrusion (apoptosis) or starting point (control) referring to apoptotic ROIs. Data shown in **G** and **H** are from n = 54 ROIs in 6 independent experiments (control or apoptosis). Scale bars, 20 µm in **A** and **C**.

### Spatial inhomogeneities in force propagation and nuclear size define which of the neighboring cells undergo cell division following apoptosis

To uncover what defines this spatial inhomogeneity of cell division, we sought out factors that are not homogeneous in space. First, the substrate deformation as measured by bead displacement is spatially inhomogeneous (Fig. 1C, 3A). We noticed that cells that underwent cell cycle progression upon apoptosis spatially correlated to the region with large displacement (Fig. 3A), indicating cells that experienced a larger displacement and consequently underwent more stretching have a higher chance to undergo cell division. Second, the spatial distribution of the size of the nucleus is spatially inhomogeneous. We found that the cells that underwent cell cycle progression upon apoptosis tended to have a larger nucleus at the beginning of apoptosis (Fig. 3B). To address whether these two spatially inhomogeneous factors, i.e., substrate deformation and nucleus size, are both required for cell proliferation, we simultaneously monitored bead displacement, nucleus size, and YAP localization. We found a case of 2 neighboring cells that both had a large nucleus (arrowhead and arrow, Fig. 3C), however only the cell located in the region that underwent large bead displacement showed YAP nuclear translocation (arrowhead, Fig. 3C). To further solidify our observation, we analyzed more than 100 cells that were in the vicinity of apoptotic cells (Fig. 3D). Only cells with a large nucleus that experienced large bead displacement showed clear YAP nuclear translocation (red, Fig. 3D). In contrast, cells with a large nucleus that experienced small bead displacement, or any cells with a small nucleus, did not show clear YAP nuclear translocation (blue in Fig. 3D). In addition, by analyzing the relationship between strain (Fig. 1K), nucleus size, and cell cycle progression measured by FUCCI, we found that only the cells with a large nucleus that experienced large strain underwent cell division (red, Fig. 3E). To rationalize how strain and nuclear size regulate YAP nuclear translocation and cell proliferation, we measured the number of nuclear pore complexes depending on nuclear size (Fig. 3F-G). The nuclear pore complex is a known structure to transport YAP from the cytoplasm to the nucleus in a tension-dependent manner [30]. We found that the number of nuclear pores increases with the size of the nucleus (Fig. 3G), and speculated that the larger nuclear size helps nuclear translocation of YAP upon mechanical stretching. Together, we found that the spatially limited cell division around the apoptotic process is defined by the spatial inhomogeneity of pre-tension release (measured by bead displacement) which leads to the strain of a cell, and the spatial inhomogeneity of nuclear size.

**Figure 3.**
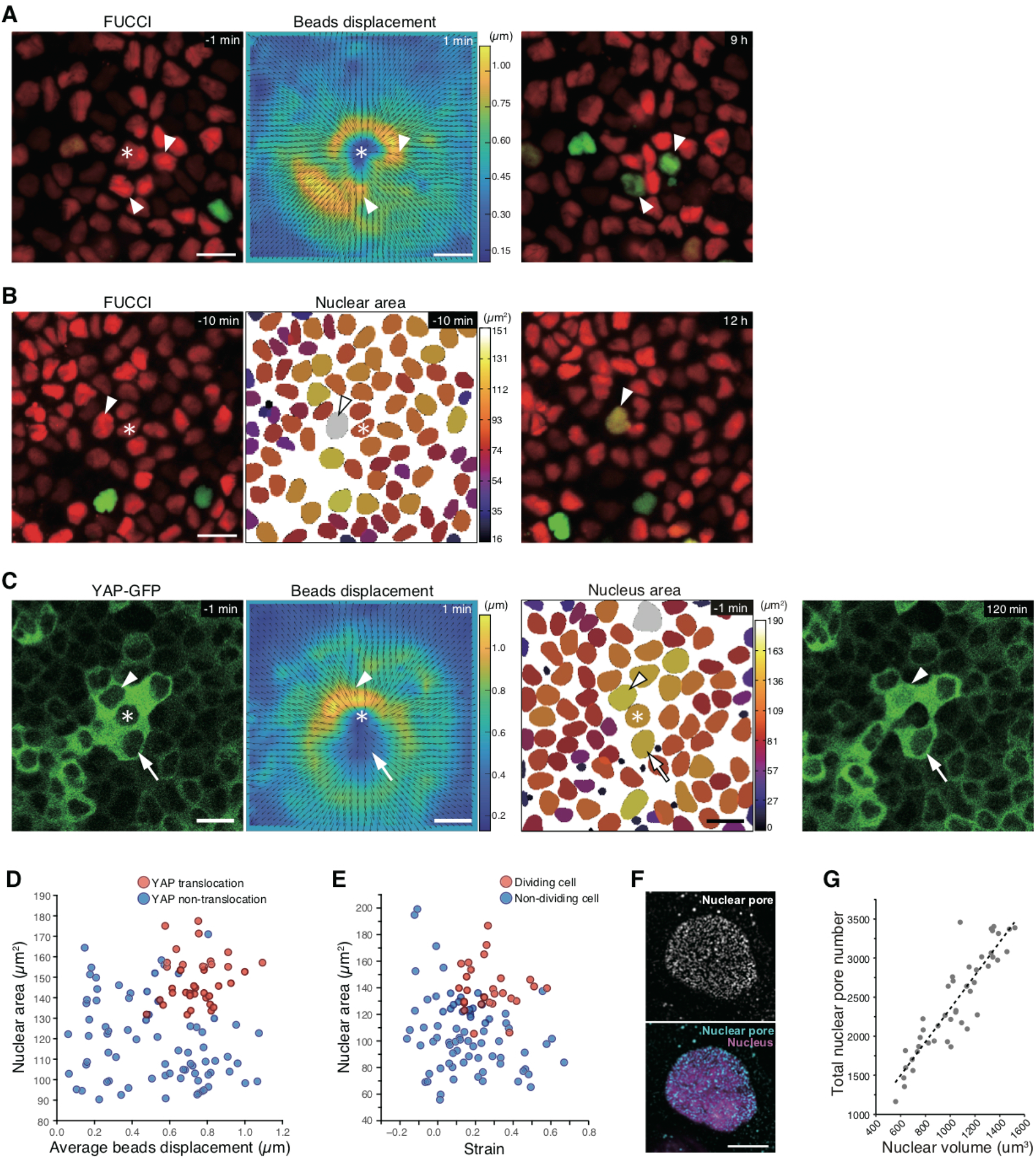
(**A** and **B**) Representative snapshots of MDCK FUCCI monolayer before (left) and after (right) apoptotic cell extrusion, and heat map with vectors of beads displacement 1 min after extrusion in **A** (middle) and heat map of the cross-sectional nuclear area 10 min before extrusion in **B** (middle). Asterisk, apoptotic cell; white arrowhead, cells underwent cell-cycle progression and subsequent cell division (data not shown); color bars, the magnitude of the displacements in µm and that of the area in µm2. **(C)** Representative snapshots of an MDCK YAP-GFP monolayer before (left) and after (right) laser induction of apoptosis, and heat map of beads displacement and nuclear area. Asterisk, apoptotic cell; white arrowhead, YAP nuclear translocation; white arrow, YAP nuclear non-translocation. **(D)** Relationship between nuclear area and average beads displacement in individual YAP-GFP+ cells for rows 1-2 at 1 min after laser induction. n = 115 cells in 36 images from 9 independent experiments. Cells were classified as YAP translocation (I_nuc/cyt_ ≥ 0.5) or non-translocation (I_nuc/cyt_ < 0.5) 2 h after laser induction. **(E)** Relationship between nuclear area and cellular strain for cells in row1 within MDCK FUCCI. For calculation of strain, cell length was measured at 0 and 30 min after cell extrusion. Cells were classified as dividing cells or non-division cells within 24 h after apoptotic cell extrusion. n = 103 cells in 16 images from 6 independent experiments. **(F)** Immunostaining for nuclear pore complex of a cell within MDCK WT monolayer. **(G)** The total number of nuclear pores is linear with respect to nuclear volume. n = 45 cells in 2 independent experiments. Scale bar, 20 µm in **A**-**C**; 10 µm in **F**.

To understand to what extent the difference in nuclear size at the beginning of apoptosis (Fig. 3D-E) represented the intrinsic inhomogeneity of nuclear size among tissue, we measured the changes in nuclear size during cell cycle progression. The nucleus size typically increased ∼10% during progression from G1 to G2 phase (Fig. S2A). Moreover, we observed a wide distribution in nuclear size within a tissue with a FWHM of 50.3μm (Fig. S2B). We thus conclude that the difference in nuclear size between cells that divided or not (Fig. 3E) and between cells that showed YAP nuclear translocation or not (Fig. 3D) did not predominantly arise from the changes in the nuclear size during cell division, but from the intrinsic inhomogeneity of nuclear size among tissue.

### Compensatory proliferation requires force propagation through the substrate

It has been shown that biochemical signaling, including mitogenic signaling, from the apoptotic cell, contributes to compensatory proliferation [3,4]. To understand to what extent the mechanical factors showed earlier play roles in compensatory proliferation in addition to mitogenic signaling, we aimed to modulate force propagation without altering the biochemical interaction between apoptotic and neighboring cells. To this end, we did not change the mechanical properties of the PDMS substrate, but altered the size of the tissue, or the friction between the gel substrate and the glass-bottom Petri dish. We first altered the size of the tissue by using microcontact printing technology to reduce the pre-tension of the tissue (Methods, Fig. 4A). The size of the mini-tissue was reduced until the bead displacement diminished. A mini-tissue of 100 μm diameter (Fig. 4A) showed a reduced bead displacement (Fig. 4B-C, Movie 6). Under this condition, the neighboring cells failed to translocate YAP into the nucleus (Fig. 4D-E, Movie 6) regardless of nucleus size (Fig. 4F) upon induction of apoptosis in a cell in the middle of a mini-tissue. Our theoretical model based on linear elasticity theory [25] predicted that the gel substrate is weakly adhered to the surface of the glass-bottom Petri dish (Fig. 1A) and that increasing this adhesion would suppress the wave propagation. To test this prediction, the dish was silanized with 3-aminopropyl trimethoxysilane (APTES) to increase adhesion of the gel to the glass (Methods, Fig. 4G). We found that under this condition the bead displacement upon apoptosis was diminished (Fig. 4H-I, Movie 7) and YAP nuclear translocation in the neighboring cells was abolished regardless of nuclear size (Fig. 4J-L, Movie 7). In addition, cell cycle progression in neighboring cells surrounding an apoptotic cell was also abolished with APTES treatment (Fig. 4M-N). Taken together, our data further support the idea that modulation of the mechanical status of the neighboring cells of the apoptotic cell through the substrate is required for the nuclear translocation of YAP, cell cycle progression, and cytokinesis of the neighboring cells.

**Figure 4.**
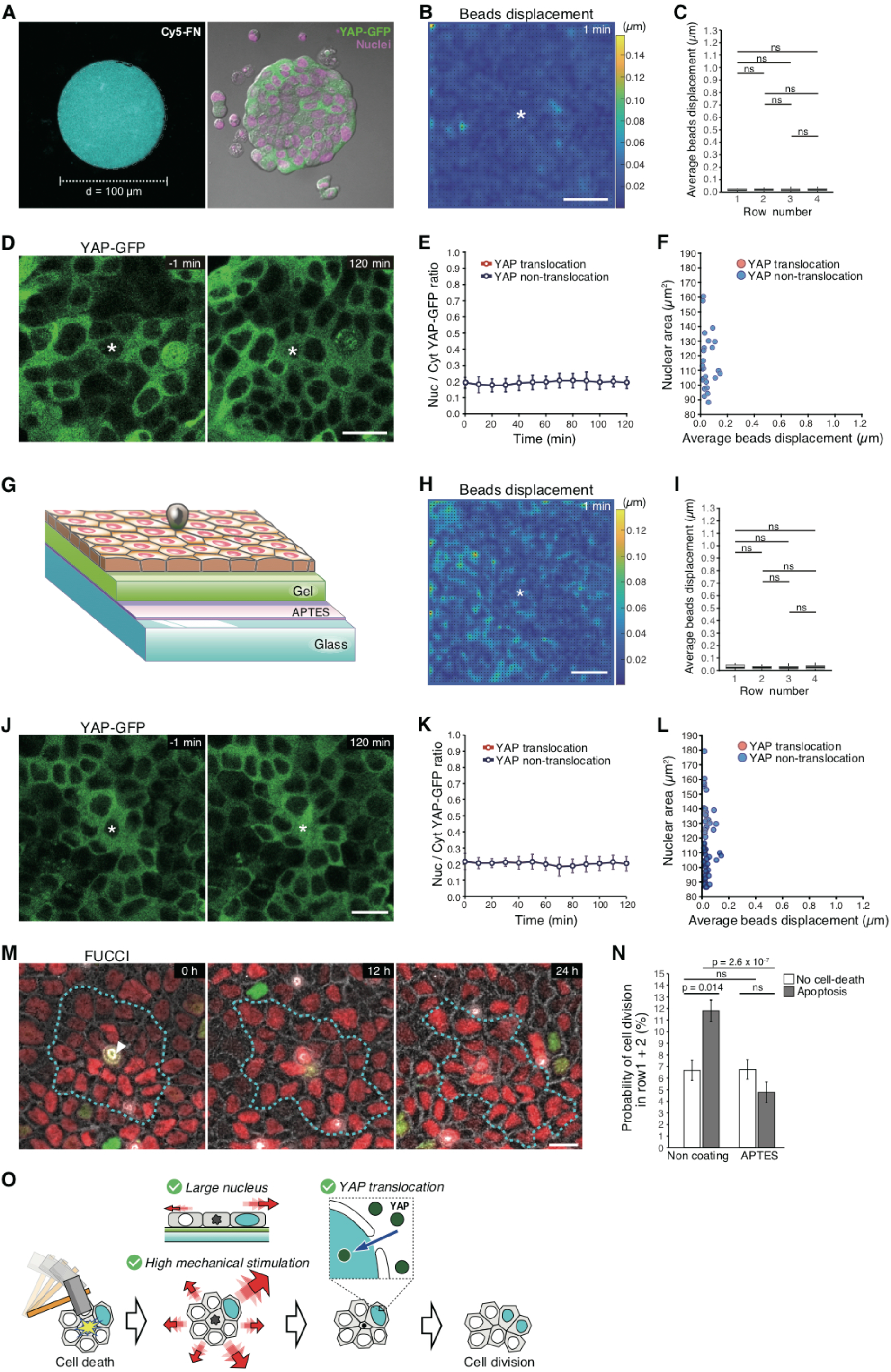
**(A)** MDCK YAP-GFP mini tissue confined on 100 µm Φ circular pattern (right). The cells are seeded on the substrate coated with Cy5 fibronectin (left, cyan). See Methods. Circular-patterned mini tissue was used in **B**-**F**. **(B)** Heat map with vectors of beads displacement 1 min after laser induction of apoptosis in **D**. Color bar, the magnitude of beads displacements in µm. **(C)** Average beads displacement for each row at 1 min after laser induction. Data are mean ± s.d., one-way ANOVA, and Levene test followed by Tukey–Kramer test. n = 5 ROIs in 5 independent experiments. **(D)** Representative snapshots of MDCK YAP-GFP cells within a confined mini tissue before and after laser induction. **(E)** Nuclear/cytosolic (Nuc/Cyt) YAP-GFP ratio in cells surrounding apoptotic cell after laser induction. n = 20 cells in 6 images from 6 independent experiments. **(F)** Relationship between nuclear area and average beads displacement in individual YAP-GFP+ cells (in rows 1-2) at 1 min after laser induction. Cells were classified as YAP translocation (I_nuc/cyt_ ≥ 0.5) or non-translocation (I_nuc/cyt_ < 0.5) 2 h after laser induction in **E** and **F**. n = 29 cells in 6 images from 6 independent experiments. **(G)** Schematic drawing of a confluent MDCK monolayer on the gel substrate which is strongly bonded to glass by APTES. APTES-coating dish overlaid with the substrate was used in **H**-**N**. **(H)** A heat map with vectors of beads displacement 1 min after laser induction in **J**. Color bar, the magnitude of beads displacements in µm. **(I)** Average beads displacement for each row at 1 min after laser induction. Data are mean ± s.d., one-way ANOVA, and Levene test followed by Tukey–Kramer test. n = 6 ROIs in 6 independent experiments. **(J)** Representative snapshots of an MDCK YAP-GFP monolayer before and after laser induction of apoptosis. **(K)** Nuclear/cytosolic (Nuc/Cyt) YAP-GFP ratio in cells surrounding apoptotic cell after laser induction. n = 23 cells in 4 images from 4 independent experiments. **(L)** Relationship between nuclear area and average beads displacement in individual YAP-GFP+ cells within rows 1-2 at 1 min after laser induction. Cells were classified as YAP translocation (I_nuc/cyt_ ≥ 0.5) or non-translocation (I_nuc/cyt_ < 0.5) 2 h after laser induction in **K** and **L**. n = 35 cells in 5 images from 5 independent experiments. **(M)** Representative time series of cells surrounding an apoptotic cell (white arrowhead) within an MDCK FUCCI monolayer. The cyan dashed line represents the boundary between row2 and row3. **(N)** Probability of cell division within 24 hours for rows 1-2 in the cases of no cell death or apoptosis under different dish conditions (non-coating vs APTES-coating). Data are mean ± s.d.; *P = 0.014 and ***P < 2.6 × 10-7, Kruskal–Wallis rank-sum test followed by Steel–Dwass test. n = 54 ROIs in 6 independent experiments (each cases with non-coating), n = 54 ROIs in 4 independent experiments (each cases with APTES-coating). **(O)** A model of compensatory cell proliferation. Asterisk in **B, D, H** and **J**, laser irradiation point; Scale bar, 20 µm.

## Discussion and conclusion

In summary, we answered a long-standing question of how only a small number of cells around an apoptotic cell undergo cell division to sustain the homeostasis of the cell number in a tissue (Fig. 4O). The spatial inhomogeneity of cell division in response to the apoptotic process is associated with inhomogeneous mechanotransduction, characterized by the nuclear translocation of YAP, and arises from the combination of the large strain and the large nuclear size of the neighboring cells. One obvious question is whether mechanotransduction in the neighboring cells is required for compensatory proliferation. Suppression of mechanical stimulus without altering the cell-cell contact between apoptotic and neighboring cells resulted in a lack of mechanotransduction and cell division (Fig. 4A-N). These results highlighted that force propagation through the substrate, which has been overlooked, plays a significant role in defining the spatial pattern of compensatory proliferation. Our theoretical model [25] suggested that the experimental condition where the gel substrate is weakly adhered to a glass-bottom dish (Fig. 1A) has mechanical features analogous to layers of tissue with viscoelasticity. We speculate that stratified epithelium, such as skin [31], shares similar mechanisms during compensatory proliferation. In contrast, the gel substrate that strongly adhered to a glass-bottom dish (Fig. 4G) behaves as an elastic substrate, which has been used for traction force microscopy [21].

Another key question is how biochemical signaling, including mitogenic signaling, that emerges from the apoptotic process, contributes to compensatory proliferation. It has been shown recently that extracellular-signal-regulated kinase (ERK) is activated in most of the neighboring cells upon apoptosis and acts as a survival factor for these cells [32,33]. Our data support the idea that mitogenic signals secreted from the apoptotic cell are crucial as demonstrated before [5–11], but not solely sufficient, to explain the spatially inhomogeneous nature of compensatory proliferation. However, we cannot rule out the possibility that there are additional mechanisms that could suppress cell division of neighboring cells next to the dividing cell. For instance, a paralog of YAP, transcriptional coactivator with a PDZ-binding motif (TAZ), is shown to be excluded from the nucleus by extrinsic compression and to cause lateral inhibition in cell fate specification [34]. Indeed, the neighboring cells that experienced a negative strain, i.e., compression, did not undergo cell division (Fig. 3E). Together, our findings complement the current biochemical understanding of apoptosis-induced compensatory growth and provide additional insights into cellular functions of how tissue precisely maintains homeostasis. Such apoptosis-induced proliferation has been found in a wide range of cancers, including breast cancer, melanoma, and glioma cells [35–38]. Our findings provide a new mechanical perspective to understand not only tissue homeostasis but also tumor pathology.

---

**Supplementary Figure 1.**
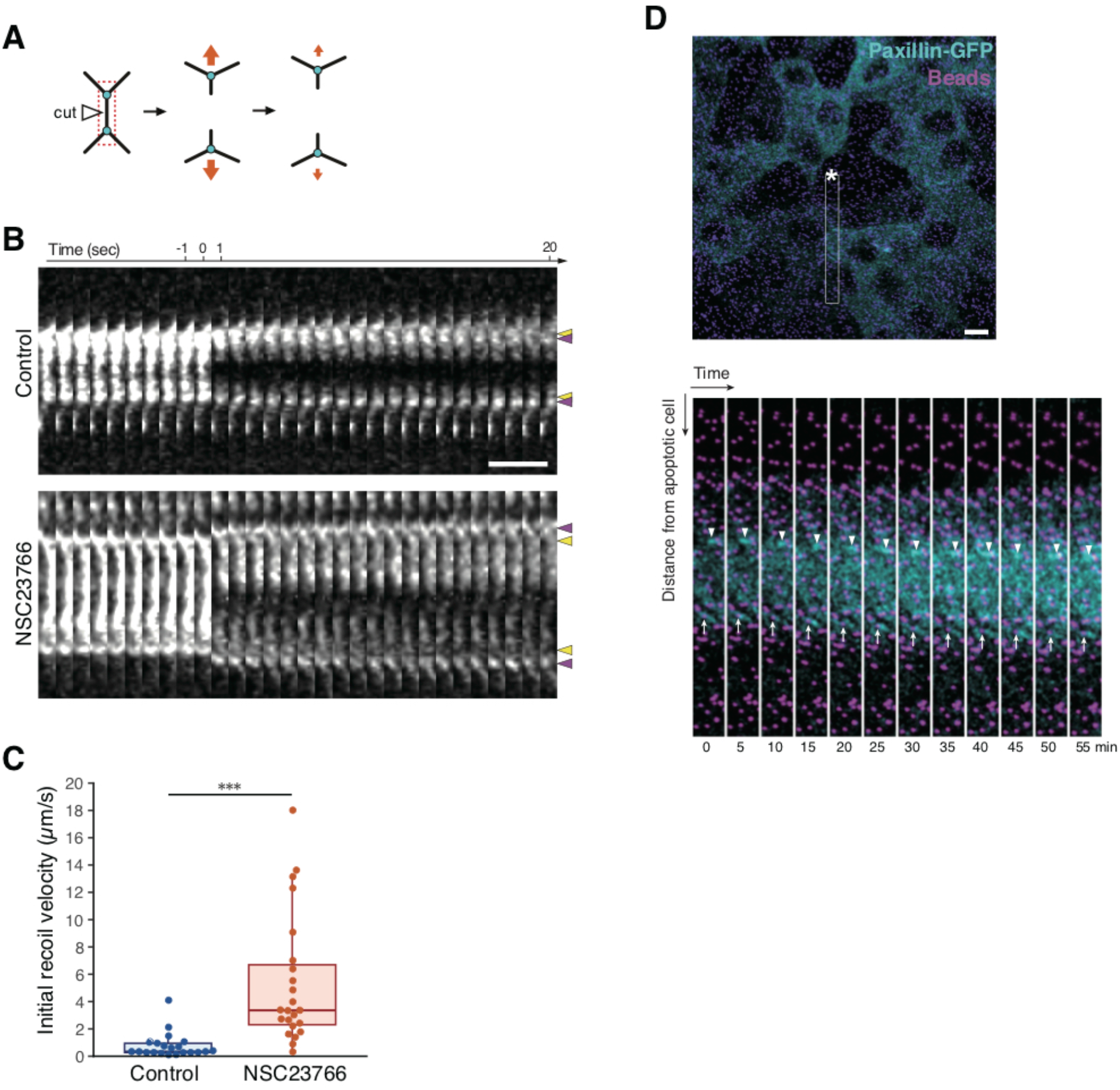
**(A)** Schematic diagram of laser ablation targeting cell-cell junction to measure the nodes (Y-junctions) displacement and calculate recoil velocity. Cyan dots, nodes of interest; white arrowhead, the point of laser ablation. **(B)** Kymographs of adherence junction labeled with E-cadherin-GFP before and after ablation, which generated from the region in the red box in **A**. Yellow arrowhead, the original positions of nodes; magenta arrowhead, the final positions of nodes after ablation. **(C)** Quantitation of initial recoil velocities after junctional ablation. Data are mean ± s.d.; ***P < 0.001, Mann–Whitney *U*-test. n = 21 ROIs (control), n = 23 ROIs (NSC23766). **(D)** Representative snapshot of MDCK Paxillin-GFP on the gel with beads (top) and the enlarged slices of movie stills enclosed in the white box after laser induction of apoptosis (bottom). Asterisk, apoptotic cell; white arrowhead, bead; white arrow, focal adhesion labeled with Paxillin-GFP. Scale bars, 10 µm in **B** and **D**.

**Supplementary Figure 2.**
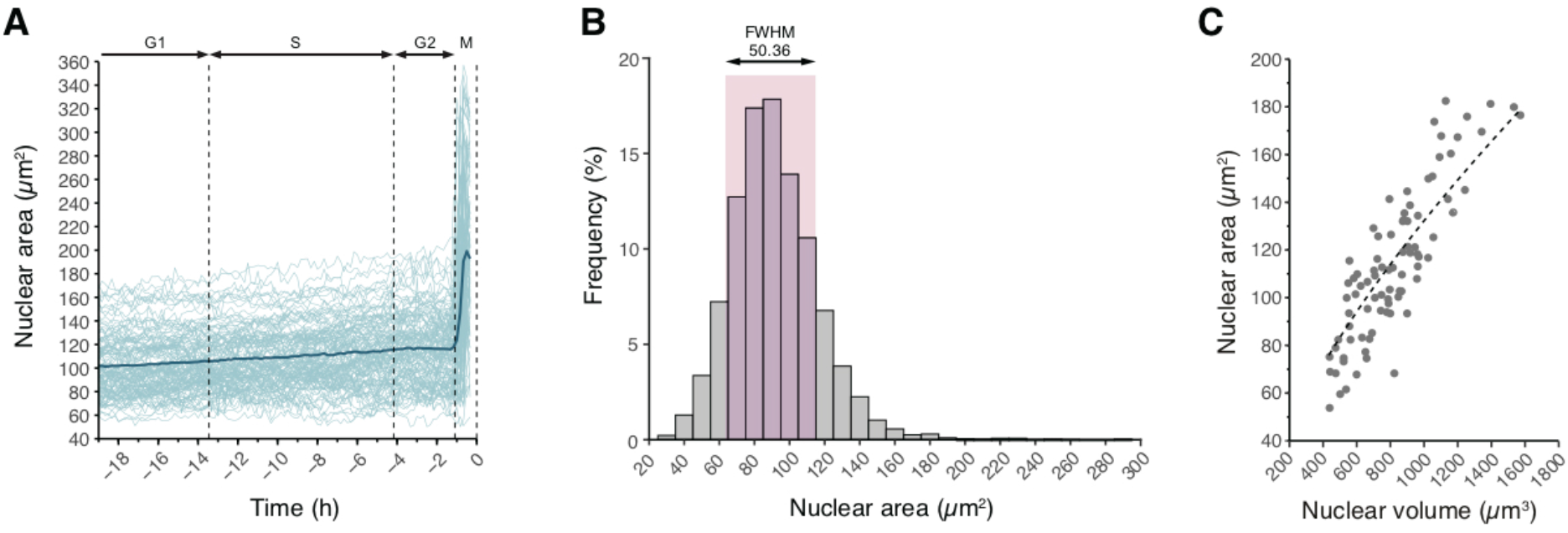
**(A)** Nuclear growth curves. Time 0 h is the time at cell division. The thick blue line represents mean nuclear area. n = 80 cells in 14 images from 4 independent experiments. **(B)** Histograms of the nuclear area within MDCK monolayers. n = 6719 cells in 10 images from 3 independent experiments. FWHM, full width at half maximum. **(C)** Relationship between nuclear area and nuclear volume of cells within MDCK monolayers. n = 86 cells in 7 images.

## Movie legends

**Movie 1**. Beads movement (left) and heat map with vectors of radial beads displacement (right) after laser induction of apoptosis within MDCK monolayer for control. The color bar indicates the magnitude of the radial outward (red) and inward (blue) displacements in µm. Scale bars, 50 µm.

**Movie 2**. Beads movement (left) and heat map with vectors of radial beads displacement (right) after laser induction of apoptosis within MDCK monolayer under NSC23766 treated condition. The color bar indicates the magnitude of the radial outward (red) and inward (blue) displacements in µm. Scale bars, 50 µm.

**Movie 3**. Tissue dynamics (DIC images) and velocity field from DIC images of MDCK monolayers after laser induction for control and NSC23766 treated condition. Red dot, laser irradiation point; length of vectors, proportional to their magnitude.

**Movie 4**. Time-lapse images of MDCK YAP-GFP monolayer stained with Hoechst 33342 (magenta). Asterisk, laser irradiation point; White arrowhead, YAP-GFP nuclear translocation; scale bars, 20 µm.

**Movie 5**. Time-lapse images of MDCK FUCCI monolayer. Black arrowhead, apoptotic cell extrusion; white arrowhead, cell proliferation; scale bars, 20 µm.

**Movie 6**. Time-lapse images of MDCK YAP-GFP mini tissue confined on 100 µm Φ circular pattern (left), beads movement (middle), and heat map with vectors of beads displacement (right). Asterisk, laser irradiation point; color bar, the magnitude of beads displacements in µm; scale bars, 20 µm.

**Movie 7**. Time-lapse images of MDCK YAP-GFP monolayer on the gel substrate which is strongly bonded to glass by APTES (left), beads movement (middle), and heat map with vectors of beads displacement (right). Asterisk, laser irradiation point; color bar, the magnitude of beads displacements in µm; scale bars, 20 µm. Note that a laser-irradiated cell and a neighboring cell underwent apoptosis.

